# Scale- and genotype-dependent spatial concordance of *Mycobacterium bovis* in cattle and badgers

**DOI:** 10.64898/2026.10.01.756065

**Authors:** Assel Akhmetova, Liliana C. M. Salvador, Katarina Oravcova, Eleanor Presho, Carl McCormick, Suzan Thompson, Lorraine Wright, Fraser Menzies, Nigel Trimble, Roland Harwood, Rowland R. Kao, Adrian Allen, Theo Pepler, Robin Skuce

**Affiliations:** School of Animal and Comparative Biomedical Sciences, University of Arizona, Tucson, AZ, United States of America; School of Biodiversity, One Health and Veterinary Medicine, University of Glasgow, Glasgow, Scotland; Veterinary Sciences Division, Agri-Food and Biosciences Institute, Stormont, Belfast, Northern Ireland; Roslin Institute and School of Physics and Astronomy, University of Edinburgh, Edinburgh, Scotland; Robertson Centre for Biostatistics, University of Glasgow, Glasgow, Scotland; Veterinary Service and Animal Health Group, Department of Agriculture, Environment, and Rural Affairs (DAERA), Belfast, Northern Ireland

**Keywords:** mammalian tuberculosis, *Mycobacterium bovis*, genetic diversity, MLVA type, spatial distribution, kernel discriminant analysis

## Abstract

*Mycobacterium bovis* causes tuberculosis (TB) in cattle, badgers (*Meles meles*), other wildlife, and humans, posing major animal and public health challenges in Britain and Ireland. Shared genotypes between cattle and badgers have been reported at regional scales, but whether this concordance holds at finer spatial scales, where control interventions operate, and whether it varies by genotype combination, remains unknown. Resolving this is critical for interpreting cross-species transmission risk and designing targeted control measures. We analyzed 1,248 *M. bovis* isolates from cattle and badgers (Northern Ireland, 1986-2018), including samples from a small, densely sampled intervention research area, using Multi-Locus Variable Number of Tandem Repeats Analysis (MLVA) and kernel discriminant analysis (KDA) across multiple spatial scales (25-95% density contours). We identified 36 MLVA types with four shared between species and one (MLV06) endemic to the area. Spatial concordance varied by type combination and scale: MLVA004-MLVA297 showed strong, bidirectional, scale-robust concordance; MLVA006-MLVA297 was concordant only at fine spatial scales; MLVA004-MLVA006 showed weak, asymmetric concordance; and no concordance was detected when all three types were analysed together. Our findings reveal spatial concordance between shared *M. bovis* genotypes varied by genotype combination and scale, indicating genotype-specific dynamics within this multi-host system.

## 1. Introduction

*Mycobacterium bovis* is the main causative agent of mammalian tuberculosis (TB) in cattle [1,2], and is transmissible within and between species, affecting domestic and wild animals, and humans [3,4]. In Northern Ireland (NI), TB remains a major animal health and economic problem, with reported herd prevalence of 14.05% for December 2025 (preceding 12 months) [5] and total annual cost of around £156 million, most of which falls on farmers, as indirect costs of around £96 million exceed direct public expenditure [6]. Human disease caused by *M. bovis* is uncommon - 702 UK notifications between 2000 and 2023, most linked to unpasteurised milk or occupational animal contact [7,8] – but NI accounts for 8.5% of these, the highest incidence of any UK region, and rates in the Republic of Ireland are among the highest in the EU [5].

Despite well-established routine cattle diagnostic testing, persistence of infection in surrounding wildlife, particularly the European badger (*Meles meles*), a well-documented reservoir host for *M. bovis* in Britain and Ireland, complicates control. Epidemiological links between cattle and badgers in NI indicate that both play important roles in transmission dynamics [9–11]. Badgers are legally protected in Britain and Ireland, and TB management has at different times and places included lethal control and vaccination of badger populations [12–14]. Reducing transmission from badgers lowers TB rates in local cattle, although comparable outcomes might be achieved through badger vaccination, fertility control, and farm movement controls [15].

Spoligotyping and multiple-locus variable number tandem-repeat analysis (MLVA) have been used for *M. bovis* molecular surveillance in NI cattle and badgers [16,17]. It has been shown that cattle-to-cattle transmission plays a key role in sustaining infection at local and national levels, likely including purchase of animals with undetected infection from other regions [18]. MLVA types from both hosts cluster spatially [19,20], and some appear region-specific across the territory [11]. However, badger roadkill surveillance in these studies was passive and analysed at province level [21], making it difficult to assess the role of badgers in *M. bovis* maintenance and transmission, or genotype distribution in the two hosts, at smaller spatial scales under systematic sampling.

A “test and vaccinate or remove” (TVR) intervention study on badgers was undertaken in NI from 2014 to 2018 in a 100 km^2^ area (plus a 2 km buffer) of County Down with high bTB prevalence and badger density (∼5.6 badger/km^2^) [22,23]. Badgers were captured and sett-side tested under field conditions using DualPath Platform (DPP) VetTB test [24], a single use lateral-flow serological assay for *M. tuberculosis* complex antibodies [25], then BCG-vaccinated and released if negative, or removed if positive (except in 2014, when test-positive individuals were vaccinated and released). Euthanised test-positive badgers underwent postmortem examination, and organ pools were cultured.

The systematic badger sampling employed in the TVR study enabled collection of a densely sampled dataset, allowing improved evaluation of associations between the distributions of *M. bovis* MLVA types in cattle and badgers at a smaller spatial scale. Specifically, we investigated: i) the genetic relatedness of MLVA types in badgers and cattle, and in cattle categories (home-bred or purchased); ii) their spatial distribution in each host; and iii) the association between *M. bovis* genetic similarity and spatial distribution between cattle and badgers.

## Materials and Methods

### 2.1 Study design

A total of 1248 *M. bovis* isolates from individual cattle and badgers were sampled in a 100 km^2^ TVR area in County Down, NI, including archival isolates to explore the relative diversity of MLVA types (Figure 1). The dataset included isolates from 1073 cattle collected within the bTB surveillance scheme during 2014-2018 and historical isolates from 1986-2013 (Figure 2); 142 isolates from 49 badgers sampled during the TVR study in 2015-2018 (in 2014 no badgers were culled, and only tracheal aspirate sampling of live badgers was conducted); and 33 isolates from 14 badgers killed in road traffic accidents (RTAs) in the TVR zone and identified as positive for *M. bovis* during 1999-2018. All isolates were associated with their year of collection and geographical coordinates.

**Figure 1.**
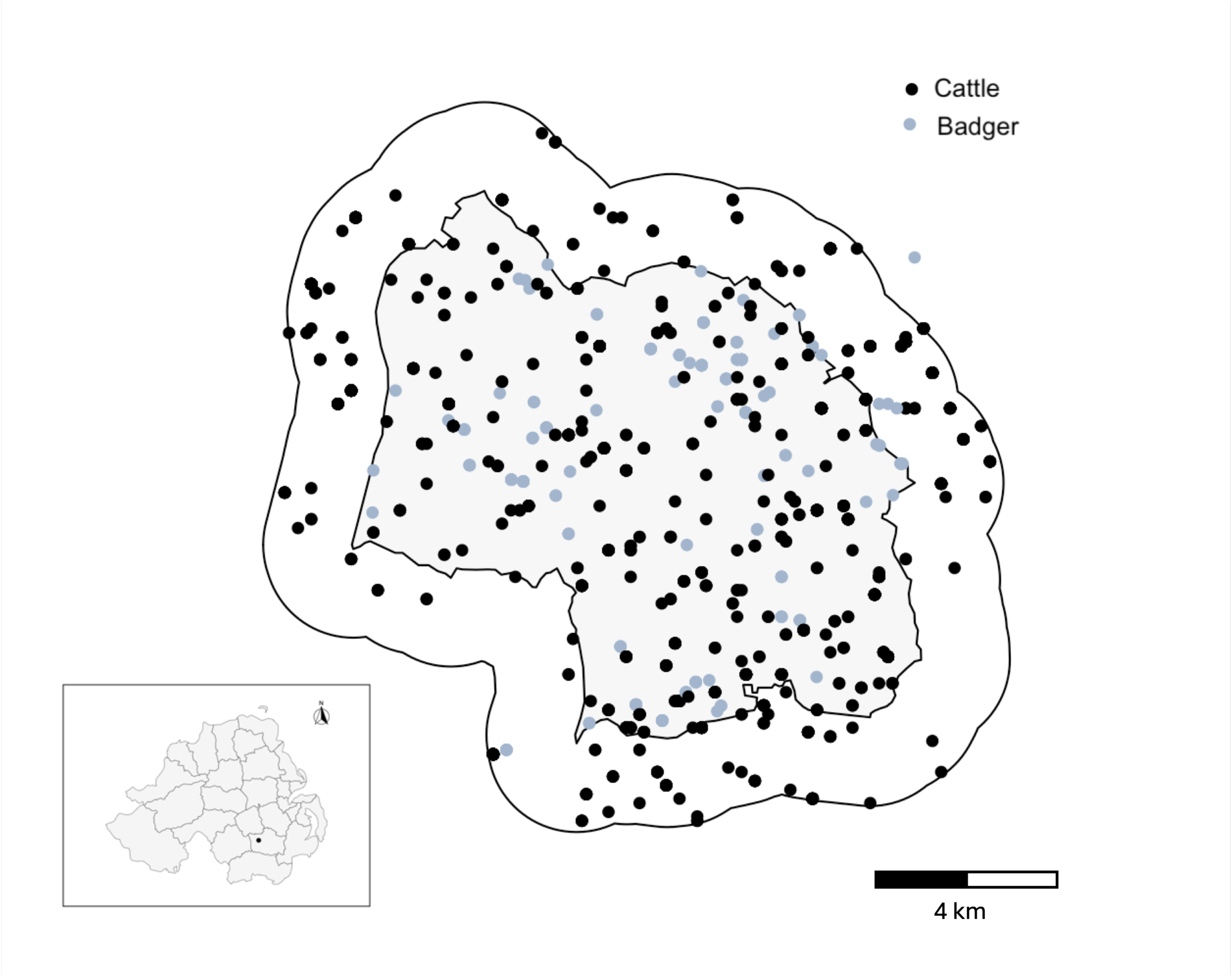
Map of study area. Map of Northern Ireland (in grey) with the TVR intervention area shown as a black point and an inset of the TVR area with zoomed locations of *Mycobacterium bovis* isolates from cattle (black dots) and badgers (light blue dots) within the TVR study area (shown in grey) and the buffer zone (shown in white). Some spatial points fall outside the buffer zone; these represent relevant historical *M. bovis* isolates that were added to the dataset.

**Figure 2.**
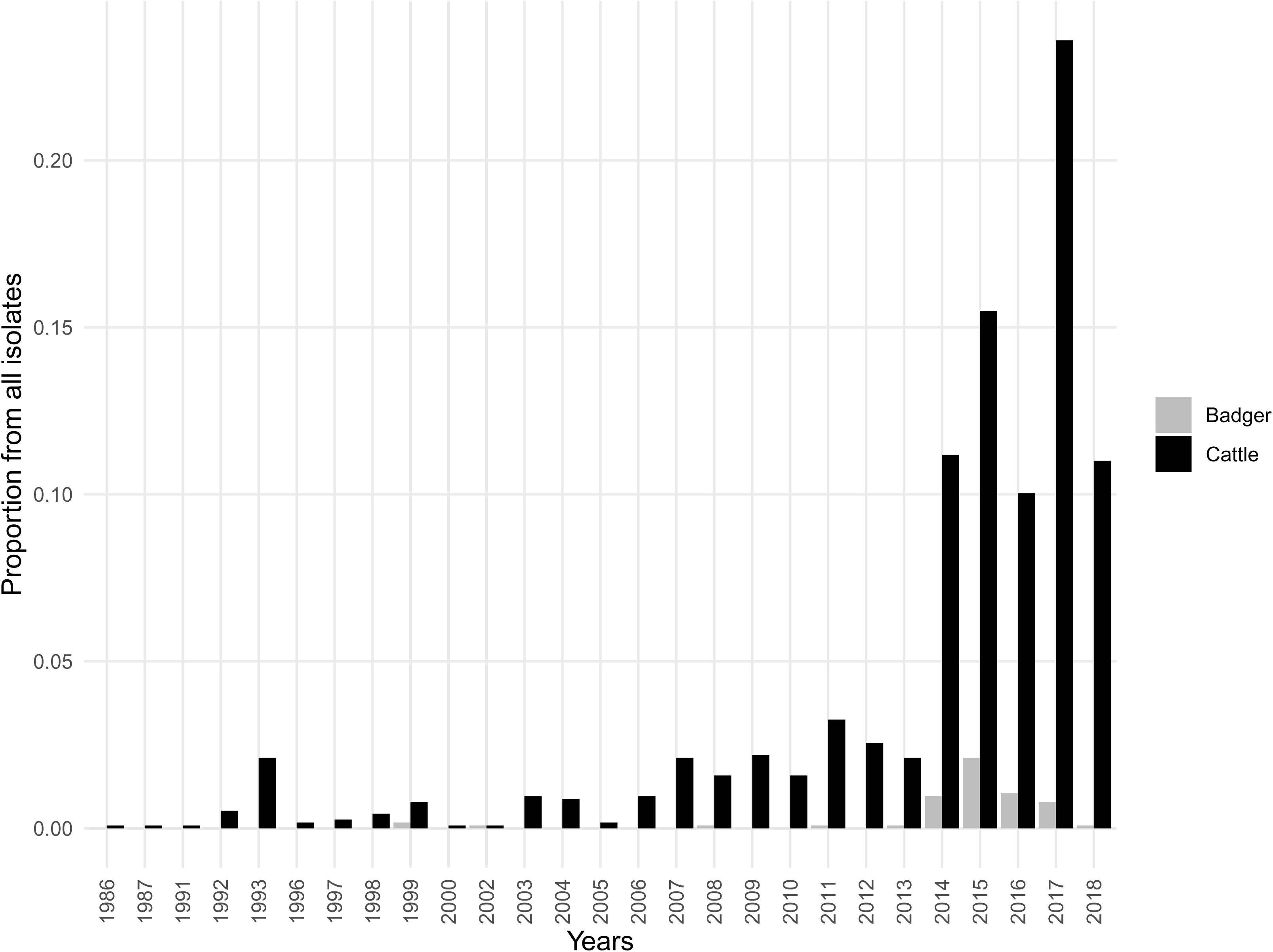
Distribution of *M. bovis* isolates by year and host-species. Distribution of *M. bovis* isolates (n = 1136) from cattle (black bars) and badgers (grey bars) by sampling years.

### 2.2 Molecular typing

For MLVA we tested eight variable number of tandem repeats (VNTR) loci: 2163b, 4052, 2461, 1955, 1895, 2165, 2163a, and 3232 [16]. This method is used for herd-and animal-level surveillance of all culture-confirmed cases in NI [16,17]; during herd-level surveillance individual cattle were identified as “home-bred” or “purchased” from official records. MLVA types were separated into groups: isolated only from badgers, only from cattle, or from both, with cattle types further discriminated as home-bred or purchased, and their distribution among these groups was assessed.

### 2.3 Genetic relatedness of M. bovis isolates

We established genetic relatedness of MLVA types using PHYLOViZ [26], a web-based tool that uses the goeBURST algorithm [27] for analysis of allelic data, providing a minimum spanning tree that groups types into clonal complexes without necessarily linking all isolates and predicts possible founding genotypes from differences in allelic profiles and number of progeny types.

### 2.4 Spatial analyses of M. bovis MLVA types

Spatial locations of individual cattle were represented by the farm buildings’ locations (available upon data sharing agreement with the data controller DAERA, NI), with individuals from the same herd “jittered” to avoid identical locations for different animals. Badger positions were trapping locations and positions of RTA badgers. All spatial points were characterized by their UK Easting and Northing coordinates. Kernel Density Estimation (KDE), using the *ks* package in R [28], was used to estimate spatial boundaries of each MLVA type found in both cattle and badgers, with at least three isolates per MLVA type per species, at density contours of 25%, 50%, 75% and 95%, the percentages representing levels of spatial density. The 95% contour, for example, delineates the area containing the 95% highest estimated density of isolates for a type.

### 2.5 Association between badger and cattle MLVA types

We used kernel discriminant analysis (KDA) [29], a nonparametric method employing KDE estimates to find spatial boundaries between the MVLA types, to investigate spatial associations between *M. bovis* genotypes in cattle and badger hosts, focusing on the three shared types (MLVA004, MLVA006, and MLVA297). KDEs were fitted for each MLVA type per host species, as described above, and comparisons between species were performed in two ways: 1) all three shared types together (MLVA004, MLVA006, and MLVA297); and 2) pairwise (MLVA004 vs MLVA006, MLVA004 vs MLVA297, MLVA006 vs MLVA297). For each analysis, we fitted a “cattle-badger” model from cattle KDEs only and used it to predict MLVA types at badger isolate locations, calculating the misclassification error rate (proportion of incorrect predictions); conversely, a “badger-cattle” model from badger KDEs only predicted types at cattle isolate locations.

To test consistency across spatial scales, all analyses were repeated restricting isolates to those within each of the four KDE contour levels (from the densest core areas, 25%, to the full distribution, 95%); analyses were performed only for types with at least three isolates per host species within a given contour level.

To assess whether observed misclassification rates reflected genuine spatial clustering versus random MLVA type distribution, we performed permutation tests with 1,000 iterations, randomly shuffling type labels within each host species while preserving spatial locations and host identity, and recalculating misclassification rates. One-sided p-values were the proportion of permuted error rates ≤ the observed rate; an observed rate below the 5^th^ percentile of the permutation distribution was taken to indicate a statistically significant association (p<0.05) between the spatial distribution of the types in the two hosts.

## Results

The availability of *M. bovis* isolates improved considerably during the TVR study period due to intensive animal-level sampling in 2015-2018. This gain was largely in badgers, as previous efforts sampled only RTA carcasses; cattle sampling remained unchanged throughout (Figure 2). Samples from 1986-2013 represent selected historical isolates from cattle gathered during routine surveillance and from RTA badgers. Isolates from 2014-2018 represent systematic sampling of all culture-positive cattle diagnosed during routine testing in the TVR area, and from 2015 onwards, all culture-positive isolates from badgers trapped within the intervention study, whether from live or post-mortem sampling.

### 3.1 Genetic diversity and relatedness of M. bovis MLVA types

The 1248 *M. bovis* isolates resolved into 36 MLVA types: 29 exclusively in cattle, three exclusively in badgers, and four (MLVA004, MLVA006, MLVA122, and MLVA297) shared (Figure 3a, b). MLVA006 was the most prevalent (51.8% of all isolates; Figure 3b) and was the focus of a phylodynamic investigation of transmission dynamics [10]. Of the 1073 isolates collected from cattle, 571 originated from “home-bred” cattle and 502 from “purchased” cattle (Figure 4). A greater diversity of MLVA types was observed in “purchased” cattle (31 types) than in “home-bred” cattle (18 types), possibly reflecting a greater diversity of source herds, whereas “home-bred” cattle showed a higher proportion of recurring types within herds.

**Figure 3.**
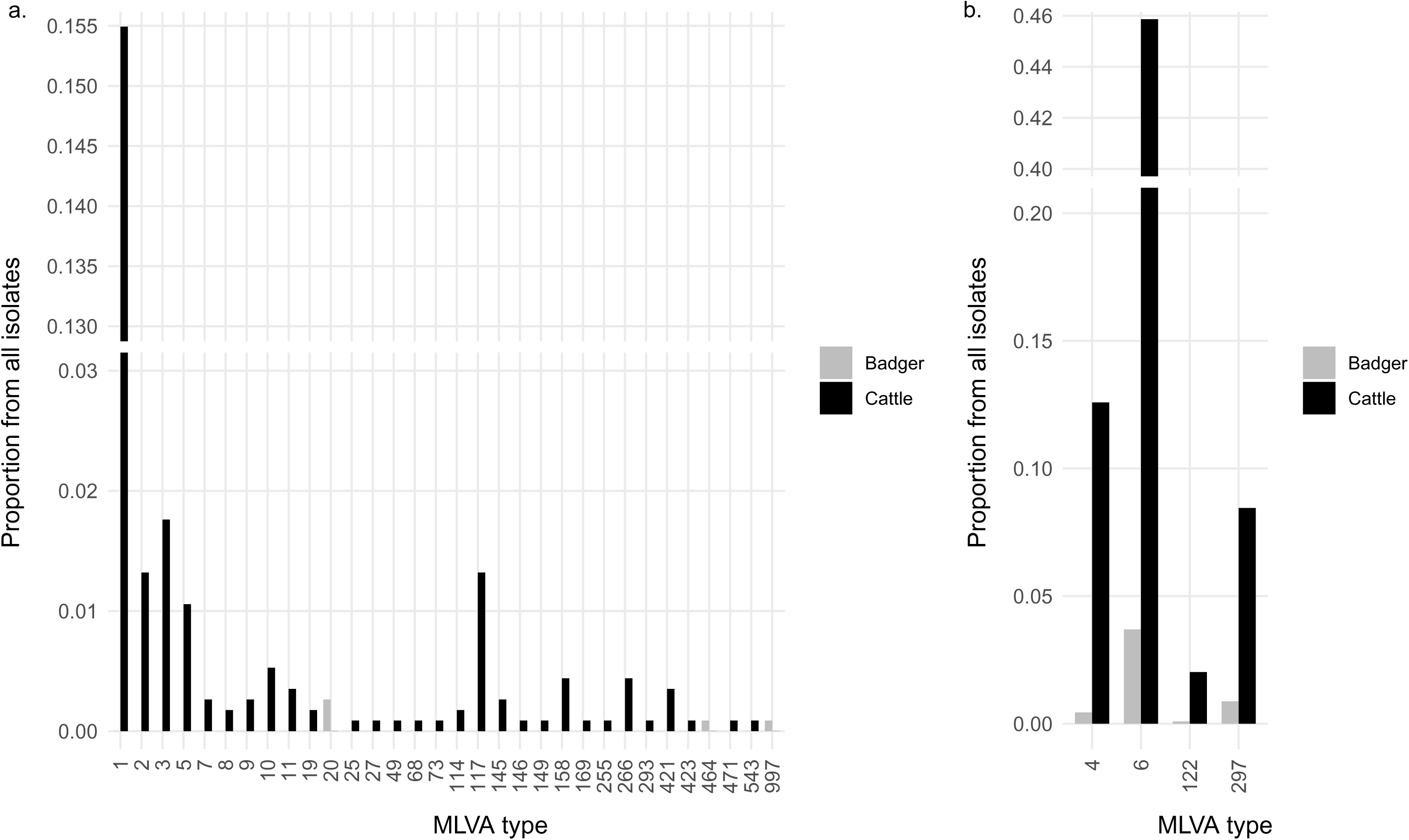
Distribution of *M. bovis* MLVA types by host-species. a. *M. bovis* MLVA types found only in cattle (black) and those found only in badgers (grey); b. *M. bovis* MLVA types shared between cattle (black) and badgers (grey).

**Figure 4.**
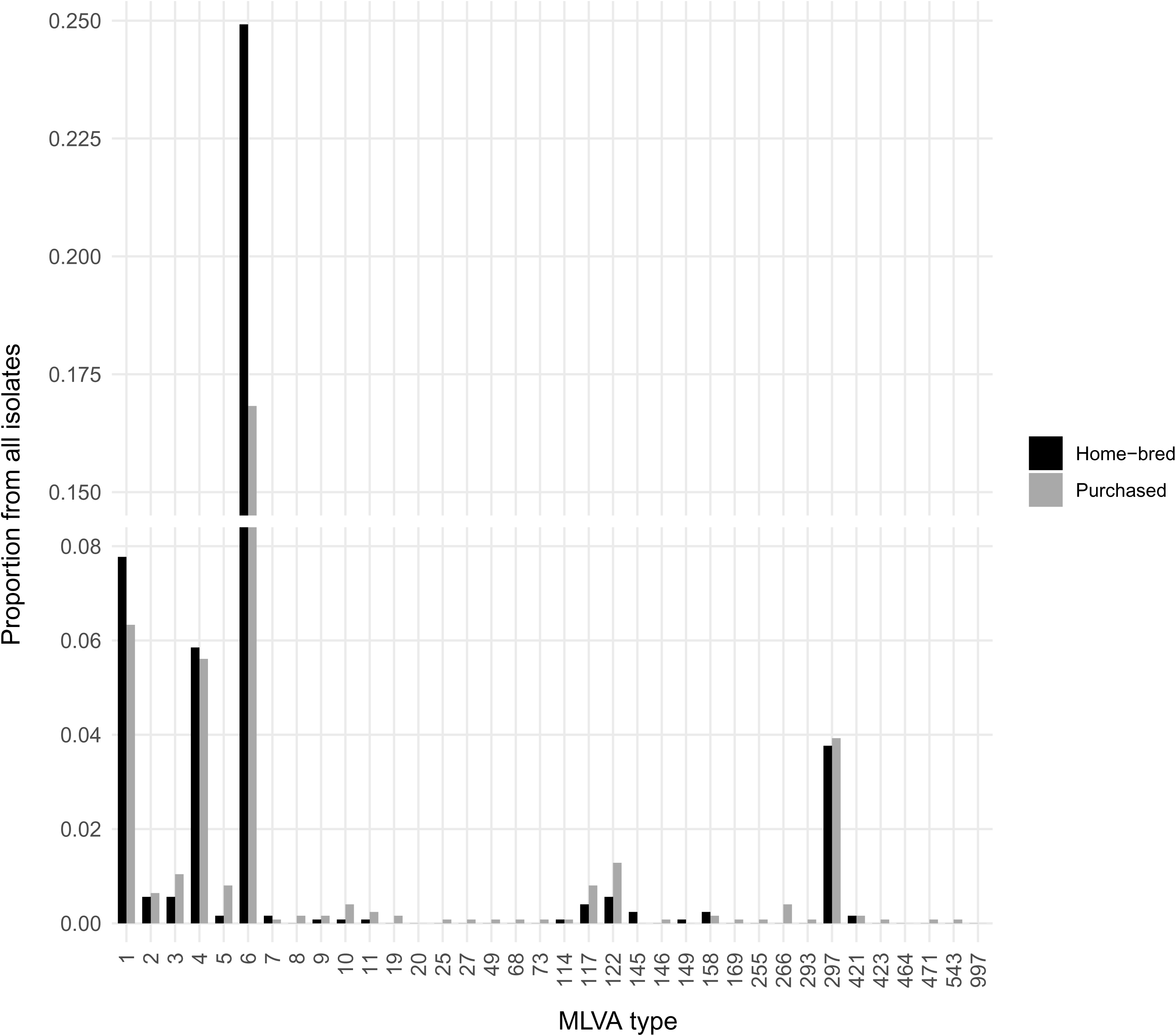
Distribution of *M. bovis* MLVA types in home-bred and purchased cattle. *M. bovis* MLVA types found only in cattle that were home-bred (black bars) and purchased (grey bars) in the area.

The minimum spanning tree derived from all MLVA types is shown in Figure 5, where the numbers on the links between nodes (in red) represent single, double and triple locus variant differences between profiles, according to the goeBURST tiebreak rules [27]. The shared MLVA006 type was the proposed founder of the clonal group and served as a central hub for the largest cluster, connecting directly to seven other types (MLVA297, MLVA421, MLVA158, MLVA464, MLVA423, MLVA266, and MLVA122) via single locus variant (SLV) links; under goeBURST rules, the node with most SLV neighbours is the most likely ancestor, and MLVA006 was also the most prevalent type and found in both hosts. MLVA001 and MLVA004 were identified as subgroup founders of their respective clusters: MLVA001, found at lower frequencies than MLVA006 and only in cattle, linked via SLV to five types (MLVA010, MLVA019, MLVA020, MLVA471, MLVA114) and to the shared MLVA004, itself central to three other types (MLVA073, MLVA005, MLVA002) via SLV links and to two other types (MLVA149, MLVA169) via double locus variant (DLV) links.

**Figure 5.**
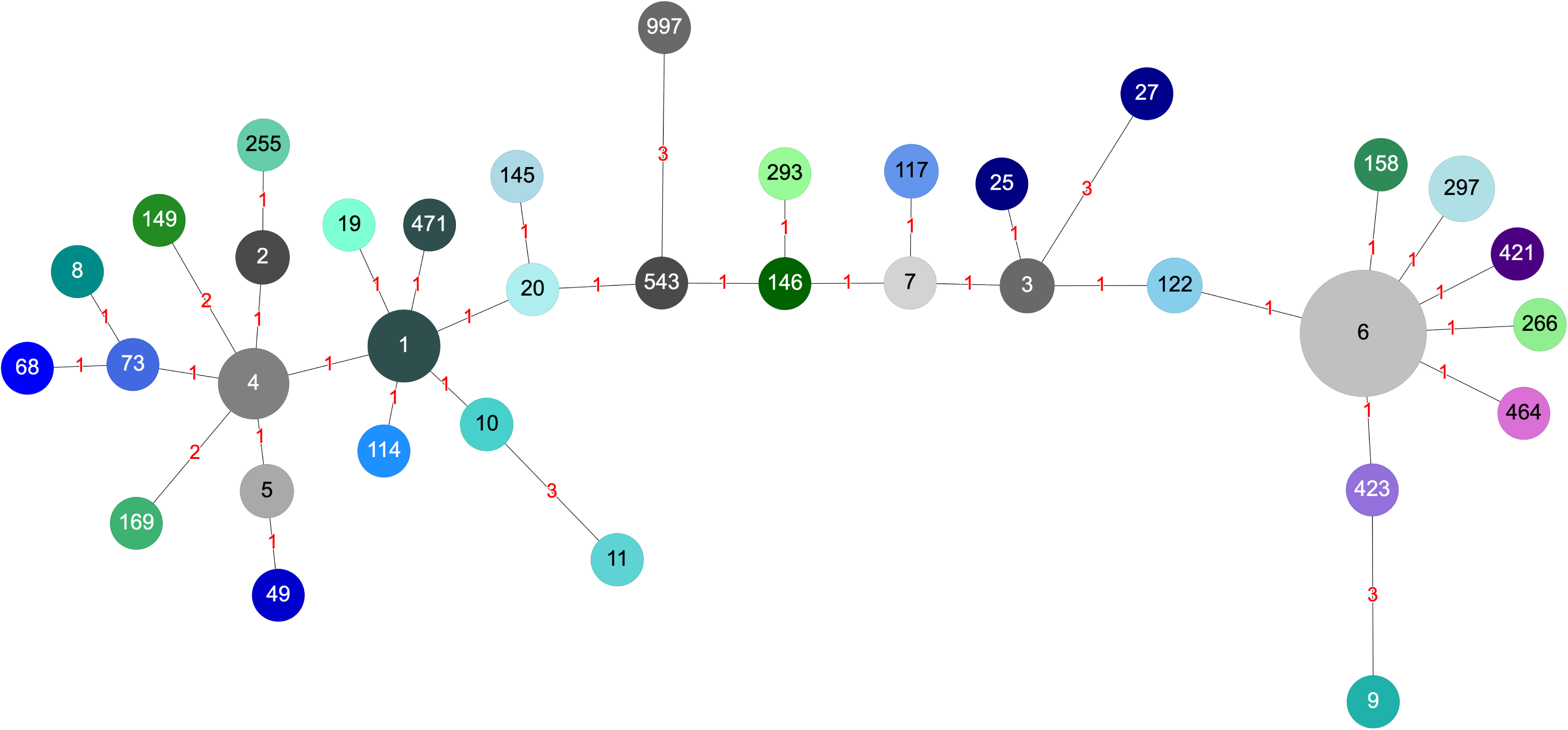
Clonal complex diagram of *M. bovis* MLVA types. The minimum spanning tree for 36 MLVA types using PHYLOViZ. MLVA006 is shown as the proposed founder of the clonal group; MLVA001 and MLVA004 are predicted subgroup founders of linked clusters. Red labels between MLVA types (nodes) on the graph demonstrate links between profiles according to tiebreak rules implemented in the goeBurst algorithm implemented in PHYLOViZ.

### 3.2 Spatial distribution of M. bovis isolates

KDE contours (Figure 6) show the core areas of highest density (25%) through the full spatial distribution (95%) for the three shared MLVA types (MLVA004, MLVA006, and MLVA297). MVLA122 was excluded for not having at least three isolates. Considering cattle and badger isolates together, the KDE boundaries of the three shared types overlap substantially across the TVR area, with MLVA006 occupying the broadest extent, consistent with it comprising the largest proportion of isolates. At 25% and 50% the contours for MLVA297 and MLVA004 are relatively distinct; at 75% and 95%, the overlap of all three types becomes more evident. The MLVA297 density estimate partially overlaps both MLVA004 and MLVA006 and covers a comparatively smaller area.

**Figure 6.**
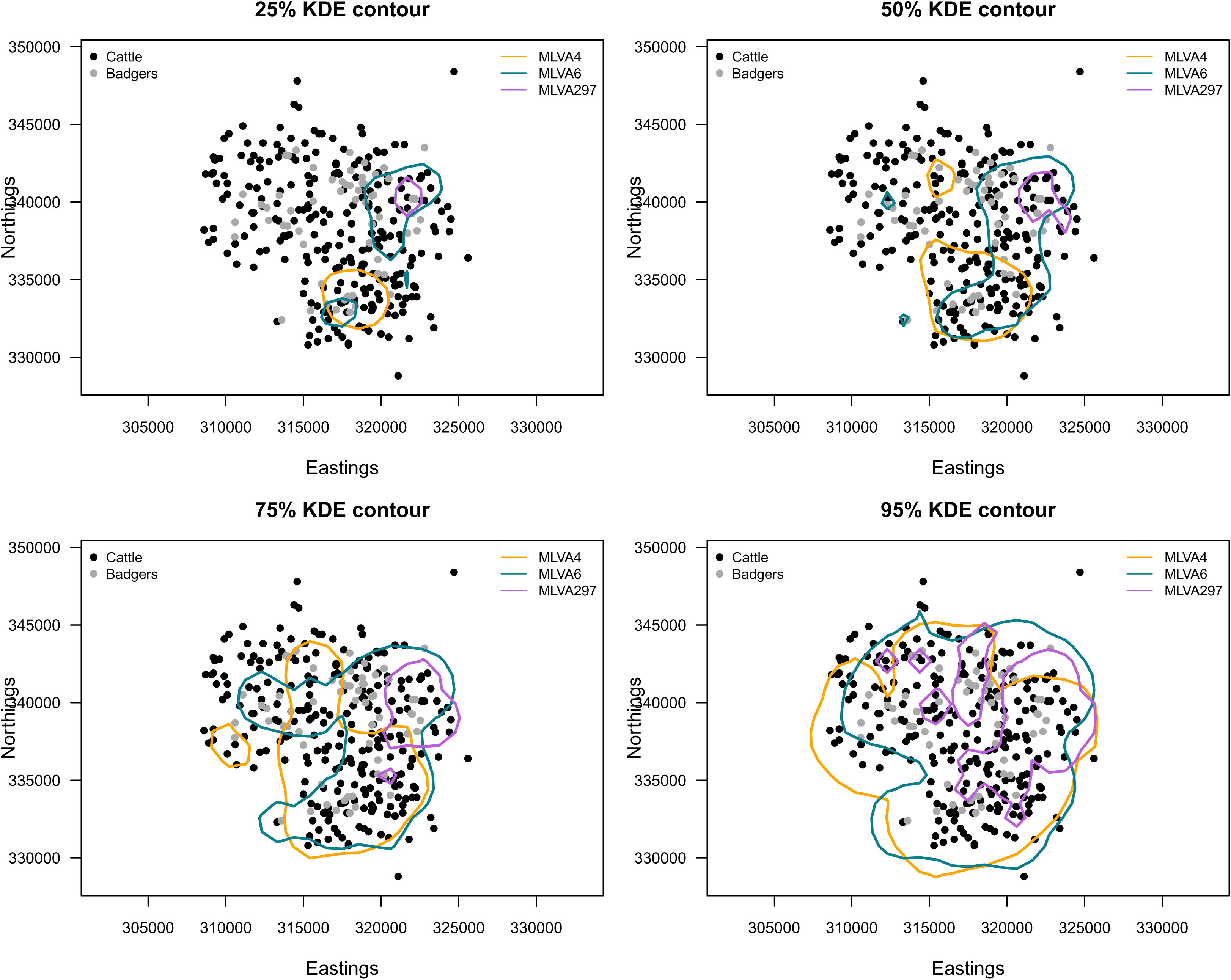
Spatial distribution of shared MLVA types at multiple kernel density levels. Kernel density estimate contours at 25%, 50%, 75% and 95% probability for the three shared *M. bovis* (MLVA004, MLVA006, and MLVA297) the three shared *M. bovis* MLVA types (orange, teal and purple lines). The geographical positions of bTB-affected farms (black dots) and culture-confirmed bTB TVR/RTA badgers (grey dots) are displayed.

### 3.3 Genetic and spatial distribution of M. bovis MLVA types between hosts

Using KDA, we tested whether the spatial distribution of shared MLVA types was concordant between badger and cattle hosts, restricting the analysis to isolates within each of the four KDE contour levels. Analyses were performed for each pairwise combination and for all shared types together, in both directions; MLVA122 and some type-KDE level combinations were excluded for not meeting the minimum of three isolates per host species. Results varied substantially across MLVA type combinations (Figure 7, Supplementary Material Table S1). MLVA004 and MLVA297 were consistently concordant in both directions at every KDE level tested (50-95%; misclassification rates 0.13-0.30, all p≤0.001), indicating that the distribution of these types in one host reliably predicted their distribution in the other, robustly across spatial scales. MLVA006 and MLVA297 showed scale-dependent concordance: significant at fine scales (25% and 50% KDE, all p≤0.008) but weakening at broader scales, particularly in the badger-cattle direction (75%: p=0.681; 95%: p=0.571), suggesting a shared spatial signal in their core area that diverges over the full distribution. MLVA004 and MLVA006 showed only weak, asymmetric concordance, reaching significance only in the badger-cattle direction at 50% and 75% KDE (p≤0.009 and 0.045), with no signal in the reverse direction, while the three shared genotypes together showed no significant concordance at any scale.

**Figure 7.**
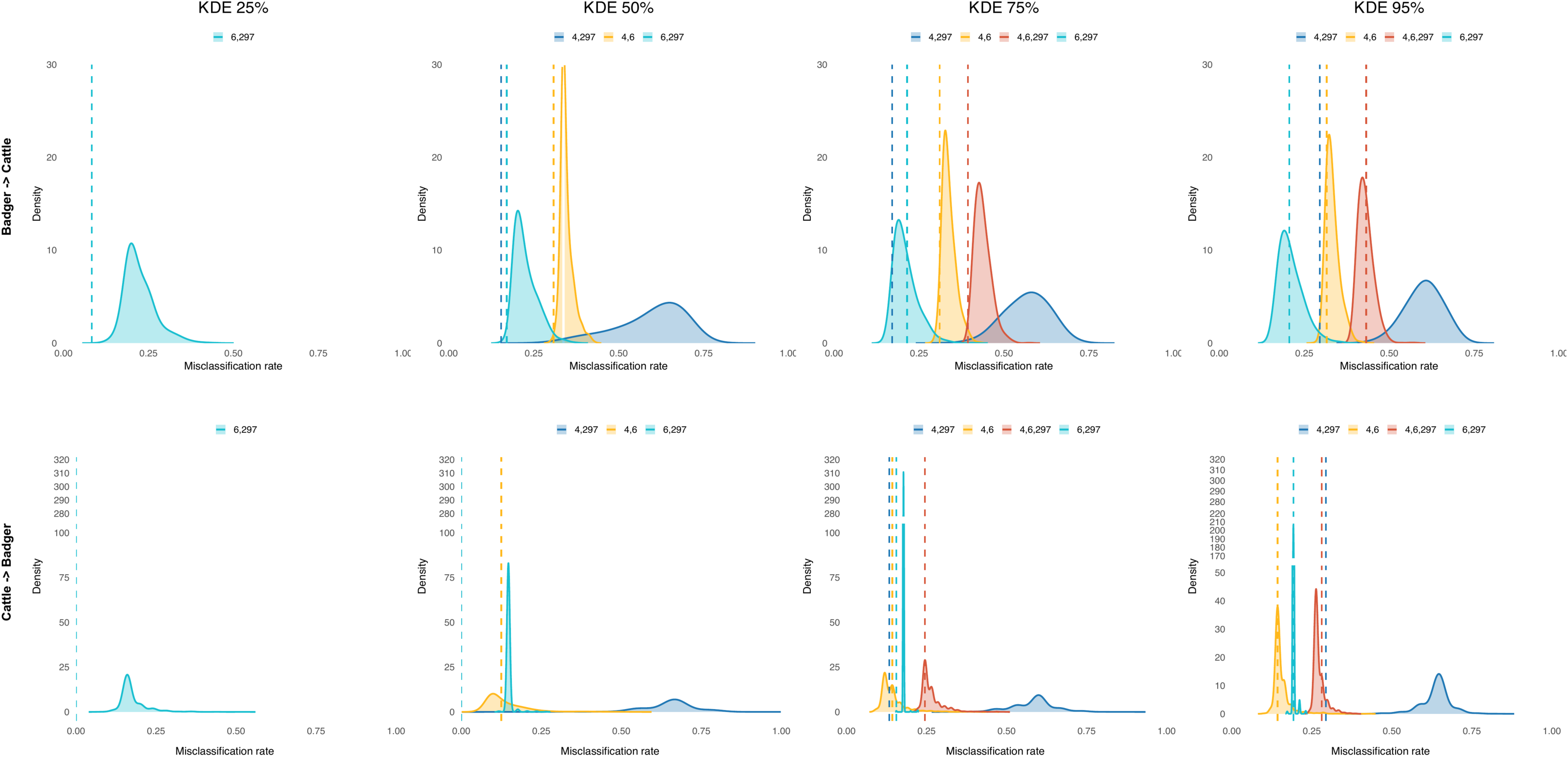
Kernel discriminant analysis for MLVA type combinations. Distribution of misclassification rates for the permutation tests for the cattle-badger and badger-cattle models (cattle KDE model predicting badger *M. bovis* MLVA types and badger KDE model predicting cattle *M. bovis* MLVA types, respectively) *M. bovis* at multiple kernel density levels (25%, 50%, 75% and 95%) for the different combinations of shared MLVA types (teal: MLVA006 and MLVA297, blue: MLVA004, MLVA297, orange: MLVA004, MLVA006, and red: MLVA004, MLVA006, MLVA297). The dotted lines represent the observed misclassification rates for each MLVA type combination.

## 4. Discussion

We investigated spatial associations between *M. bovis* MLVA types in cattle and badgers using a large dataset collected mainly during the TVR intervention study, with intensive, animal-level, systematic sampling in a 100 km^2^ area. Previous studies showed most types to be geographically clustered, suggesting transmission between cattle and badgers at a regional scale [30], but badger sampling relied on RTA surveillance, at lower numbers and resolution, so a direct quantitative relationship could not be determined. The goeBurst analysis [27] shows the main MLVA types and their daughter types to be closely related: MLVA006, MLVA122 and MLVA297 are endemic to the TVR area (Figure 5), MLVA297 being less frequent but spatially restricted (Figure 6). MLVA004 and MLVA006 were common in cattle surveyed at the herd level in 2003-2016 and co-localised in RTA badgers [30]; MLVA006 is strongly localised to the Newry Divisional Veterinary Office (DVO) region [30], which includes the TVR site (Figure 1), explaining its high isolation frequency and that of its progeny here (Figure 5). MLVA004 was found in only a few badger cases, consistent with survey data showing it is not endemic to the TVR area but was recently introduced from elsewhere in NI, most likely via cattle movement, with spillover to badgers [11]. Several other types fell outside the main clonal complex or differed at more than three of the eight loci (MLVA009, MLVA011, MLVA027, MLVA149, MLVA169, and MLVA997), some isolated only once; their rarity may reflect infection introduced rather than circulating locally, though our data cannot distinguish introduction via purchased cattle, transient undetected cattle infection, or badgers ranging in. The kernel density contours of the three shared types overlapped from KDE 50% onwards, MLVA006 and MLVA297 already overlapping at the 25% (core density) contour. MLVA297 showed the slowest boundary expansion and the most restricted distribution, consistent with higher local persistence; overlap of all three at KDE≥75% demonstrates that at least three different MLVA types coexist and co-circulate across the study area in both hosts. Notably, the MLVA297 kernels were almost entirely nested within the broader MLVA006 kernels at every contour level (Figure 6), consistent with local descent of MLVA297 from MLVA006, daughter types emerging from a dominant parental strain, as reported previously [19].

Using density estimates for shared types, *M. bovis* presence in cattle can predict its presence in badgers and vice-versa, but the strength, direction, and scale of this signal varied across types. For MLVA004 and MLVA297 prediction was symmetric and robust across all scales tested (50-95% KDE), indicating stable, mutually informative concordance. Symmetry matters: one-directional significance can arise from differences in sample sizes or spatial coverage, whereas significance in both directions suggests a shared spatial process acting on both hosts. No consistent signal was detected for MLVA006 with MLVA004, likely because of the endemic status and high local frequency of MLVA006 - herd-level NI surveillance 2003-2016 recorded it at 8.56%, highly localised to Newry DVO [17], in which the TVR zone is nested - so its distribution is diffuse and predictive models based on it become uninformative; alternatively, this absence may indicate a dynamic, spreading process rather than a settled endemic distribution. The scale-dependent concordance between MVLA006 and MLVA297 is complementary: significant concordance in core density areas that weakens at broader scales is consistent with a locally restricted sub-lineage embedded within the distribution of its parent strain; MLVA297 is genetically close (one locus apart). Applied across multiple contour levels, the KDA framework may help detect geographically restricted sub-lineages of endemic strains, while the scale-robust association between MLVA004 and MLVA297 illustrates its potential for newly introduced types: because not all types were found in wildlife, most imported types apparently did not persist in the TVR zone, whereas MLVA004 did - incursions may be common, but few establish successfully. It may also help localise spillover, since badgers are highly philopatric [31] and TVR badgers, sampled at sett sides, have more precise spatial data. Importantly, these patterns are spatial signatures consistent with cross-species transmission rather than direct evidence of it, since shared environmental exposure to type-specific strain pools could produce similar patterns; whole-genome sequencing (WGS) of paired badger-cattle would resolve sequence types and help distinguish transmission mechanisms, whereas MLVA with this spatial approach is a simple, inexpensive screen for locally established incursions to flag for WGS.

In the TVR study, badgers testing DPP-positive dropped from 14% (95% CI 0.1-0.20) in 2014 to 1.9% (95% CI 0.8-3.8) in 2018 [32]. Studies in the UK and Republic of Ireland demonstrated significant bTB reductions in proactive, non-selective badger culling area [30], although surrounding reactive and no-culling areas showed a negative effect, potentially due to badger perturbation [33]; TVR instead used sett-side DPP serology to guide selective removal alongside BCG vaccination of test-negative badgers [23]. While our earlier pathogen genomics research suggested that cattle-to-cattle transmission dominates here [10,11], badgers still contribute to maintenance and transmission, if at a lower level than cattle [10], and even a minor direct role can seed outbreaks subsequently amplified within herds [12]. Because badgers are philopatric [31], the *M. bovis* they introduce recurs in the same places, sustaining infection within the badger-cattle complex.

Several limitations apply. The badger data combined pre-TVR RTA cases with systematically sampled TVR data [10,11,23], so changes in the kernel densities of the shared types over time may not be accounted for. MLVA has lower discriminatory power than WGS [34–36], making genotype relatedness difficult to assess and tending to overestimate genetic differences between types. *M. bovis* has a clonal structure, evolves slowly, and isolates from the same area tend to be closely related genomically [37]. We did not consider control measures, herd types, and explicit animal movements [17,30], and cattle locations, jittered around the farm center, were less precise than badger locations, while land fragmentation [38] and within-farm features may influence cattle cluster positions.

Previous studies suggested cattle play a major role in *M. bovis* transmission in NI [10,39], but earlier badger data collection was less systematic than during the TVR. Our findings provide further evidence of this at smaller spatial scales. With more intensive isolation from badgers, and despite greater MLVA diversity in cattle, we show co-circulation of shared types, presence in one host predicting the same genotypes in the other. Although the model does not indicate the direction of transmission, incorporating these data into epidemiological models is fundamental to understanding transmission dynamics in multi-host systems. In conclusion, despite lower resolution, MLVA genotyping is a robust, useful technique for rapidly identifying and targeting outbreaks amenable to higher resolution investigation and phylodynamic analyses using WGS.

## Acknowledgements

The authors thank DAERA Veterinary Service and Animal Health Group staff involved in collecting and processing data from cattle and badgers, and AFBI Veterinary Sciences Division staff for generating the *M. bovis* MLVA types. During the preparation of this work, the authors used an AI language model (Claude Opus 5.5 Anthropic) to assist with condensing the text to meet the journal’s word limit. The authors reviewed and edited all output and take full responsibility for the content of the publication.

## Author Contributions

Conceptualization: FDM, RRK, AAl, RS; Data curation: EB, CM, ST, LW; Formal analysis: AAk, LCMS, TP; Funding acquisition: FDM, RRK, AAl, RS; Investigation: AAk, LCMS, EB, CM, ST, LW, AAl, RS; Methodology: AAk, LCMS, TP; Project administration: NT, RH, AAl, RS; Supervision: LCMS, KO, RRK, TP; Writing – original draft: AAk, LCMS, TP; Writing – review & editing: AAk, LCMS, KO, RRK, AAl, TP, RS.

## Data Accessibility

Data used in this study are presented in Supplementary Material Table S2. Cattle herd information such as herd ID, number of animals per herd, and spatial location data are anonymized and are protected under a data sharing agreement. The R scripts used for this publication are freely available on the Github link below: https://github.com/lsalvador/mbovis-mlva-spatial-concordance.

