## Supplementary Materials for "Scale- and genotype-dependent spatial concordance of *Mycobacterium bovis* in cattle and badgers"

### **Affiliations**

**Supplementary Tables**

**Table S1. Results of Kernel Discriminant Analysis.** Combination: combination of MLVA types to be analyzed; kde_pct: kernel density estimate percentage (25%, 50%, 75%, 95%) used for the KDA analysis; n_badgers_total: number of badgers for each pair combination analysis; n_cattle_total: number of cattle for each pair combination analysis; n_badgers_by_genotype: number of badgers by genotype; n_cattle_by_genotype: number of cattle by genotype; rate_badger_to_cattle: KDA misclassification rate from badgers to cattle; pvalue_badger_to_cattle: statistical support for KDA misclassification rate from badgers to cattle; rate_cattle_to_badgers: KDA misclassification rate from cattle to badgers; pvalue_cattle_to_badger: statistical support for KDA misclassification rate from cattle to badgers.

**Table S2. Badger and cattle metadata used in this study.** No.: sample ID; Survey: TVR: part of the TVR area; DVO: Species (either badger or cattle), cattle are represented by zone (TVR zone) or 2km (2 km buffer zone around TVR area); Year: year of sample collection; isolate per badger: number of isolates per badger; RTA: badger samples collected after a road traffic accident (RTA); Herd ID: ID of cattle herd (anonymized); animal per herd: number of animals per herd (anonymized); H/B: indication if the cattle in focus were homebred (H) or bought in (B); String: eight variable number of tandem repeats (VNTR) that were tested; Genotype: Multi-Locus Variable Number of Tandem Repeats Analysis ID; Gen+Spol: MLVA type + spoligotype; x coord: badgers: x coordinate of the position of where the badger was found, cattle: x coordinate of farm’s location (anonymized); y coord: badgers: y coordinate of the position of where the badger was found, cattle: y coordinate of farm’s location (anonymized).
